# The ancestral chromatin landscape of land plants

**DOI:** 10.1101/2022.10.21.513199

**Authors:** Tetsuya Hisanaga, Shuangyang Wu, Peter Schafran, Elin Axelsson, Svetlana Akimcheva, Liam Dolan, Fay-Wei Li, Frédéric Berger

## Abstract

**Background:** In animals and flowering plants specific chromatin modifications define three chromosomal domains: euchromatin comprising transcribed genes, facultative heterochromatin comprising repressed genes, and constitutive heterochromatin comprising transposons. However, recent studies have shown that the correlation between chromatin modifications and transcription vary among different eukaryotic organisms including mosses and liverworts that differ from one another. Mosses and liverworts diverged from hornworts, altogether forming the lineage of bryophytes that shared a common ancestor with all land plants. We aimed to obtain chromatin landscapes in hornworts to establish synapomorphies across bryophytes.

**Results:** We mapped the chromatin landscape of the model hornwort *Anthoceros agrestis.* By comparing chromatin landscapes across bryophytes we defined the common chromatin landscape of the ancestor of extant bryophytes. In this group, constitutive heterochromatin was characterized by a scattered distribution across autosomes, which contrasted with the dense compartments of heterochromatin surrounding the centromeres in flowering plants. Topologically associated domains were primarily occupied by transposons with genes at their boundaries and nearly half of the hornwort transposons were associated with facultative heterochromatin and euchromatin.

**Conclusions:** Most of the features observed in hornworts are also present in liverworts but are distinct from flowering plants. Hence, the ancestral genome of bryophytes was likely a patchwork of units of euchromatin interspersed within facultative and constitutive heterochromatin and each unit contained both transposons and genes sharing the same chromatin state. We propose this genome organization was ancestral to land plants and prevented transposons from being segregated as constitutive heterochromatin around point centromeres as in flowering plants.

## Background

A hallmark of eukaryotes is the association of their genome with nucleosomes composed of 147 bp of DNA wrapped by two heterodimers of histones H2A and H2B and a tetramer of histone H3 and H4 [1–4]. A plethora of post-translational modifications (PTMs) on N- and C-terminal tails of core histones have been identified including methylation, phosphorylation, and acetylation of H2A, H2B, and H3 that are conserved and found in similar genomic contexts across eukaryotes [5–8].

PTMs signal the position of the transcriptional start sites, gene bodies and terminators [9–13] and are involved in the regulation of cell cycle checkpoints, heterochromatic formation, centromere assembly, DNA replication, DNA repair, and gene transcription among others [14,15].

In euchromatin, trimethylation of lysine 4 on histone H3 (H3K4me3) is a PTM associated with the transcription start sites (TSS) of transcribed genes [9,16] while H3K36me3 is associated with transcriptional elongation [17]. Facultative heterochromatin forms nuclear domains marked by H3K27me3, which is deposited by Polycomb Repressive Complex 2 (PRC2) [18,19]. H3K27me3 is crucial for silencing developmental genes [20–22]. By contrast, constitutive heterochromatin is occupied by transposons and is marked by H3K9 methylation in yeast, animals, and flowering plants [23]. In yeast and animals, H3K9me3 is bound by heterochromatin protein 1 (HP1) which promotes compaction of heterochromatin and forms a compartment that recruits other heterochromatin interacting proteins [24]. In flowering plants, H3K9me2 does not bind to HP1 but instead recruits the plant-specific DNA methyltransferases CHROMOMETHYLASE (CMT) 2 and 3 [13]. CMT2/3 deposits methyl groups specifically on cytosines in the CHG context [25]. H3K9me2 also recruits the *de novo* DNA methyltransferases DOMAINS REARRANGED METHYLASE 1(DRM1) and DRM2[26] and DNA methylation is bound by a domain present in the SU(VAR)3-9 HOMOLOG (SUVH) 4/5/6 that methylates H3K9 [26]. Hence, feed-back loops involving H3K9 methylation and DNA methylation maintain heterochromatin in flowering plants. In addition, H3K27me1 is deposited on the histone variant H3.1 at constitutive heterochromatin of *Arabidopsis thaliana* by the specific histone methyltransferases ARABIDOPSIS TRITHORAX-RELATED 5 (ATXR5) and ATXR6 [27].

Although histones PTMs and their function are conserved amongst angiosperms, recent investigations have begun to show a fairly distinct chromatin landscape in bryophytes, which diverged from vascular plants ca. 500-480 MYA. Bryophytes comprise three monophyletic groups – hornworts, liverworts, and mosses [28]. Five histone PTMs and DNA methylation have been profiled in the genomes of two model bryophytes, the moss *Physcomitrium patens* [29,30] and the liverwort *Marchantia polymorpha* [31,32]. DNA methylation in the CG context is maintained by an ortholog of the flowering plant METHYLTRANSFERASE 1 (MET1), but other pathways that control DNA methylation in angiosperms are not conserved in bryophytes. In *P. patens*, de novo DNA methylation depends on DNMT3, which is distinct from DRMs involved in the RNA-directed DNA methylation (RdDM) pathway in *A. thaliana* [29]*. M. polymorpha* has two CMT-like proteins which reside outside of the clade containing CMT3 [33]. In addition, horizontal gene transfer of a prokaryotic N4 methyltransferase led to N4 methylation in sperm of *M. polymorpha* [34]. While the enzymes that deposit H3K9 methylation have not been characterized in bryophytes, orthologs of SUVH 4/5/6, which are responsible for the deposition of this PTM in *A. thaliana*, are present in bryophytes [35]. H3K27me1 is also present at heterochromatin in *M. polymorpha* [32]. While H3K36me3 marks the body of actively transcribed genes in *M. polymorpha* and *P. patens*, as it does in *A. thaliana*, H3K4me3 appears to be associated with repressed genes in *M. polymorpha* but not in *P. patens* [30,32]. Forty percent of transposons are covered by H3K27me3 in *M. polymorpha* [32], transposons in *P. patens* are mostly covered by H3K9me2 [30]. Thus, *M. polymorpha* and *P. patens* may differ in the mechanisms associated with DNA methylation and PTMs and their relationship with transcriptional states, genes, and transposons and it is not possible to draw an overview of the chromatin landscape common to bryophytes based on these representatives of liverworts and mosses, respectively.

Hornworts form a monophyletic group that diverged before the divergence of liverworts and mosses and thus the traits common to hornworts and mosses or liverworts are hypothetically representative of the ancestral traits of bryophytes [28,36]. To reconstruct the ancestral chromatin landscape of bryophytes, we investigated DNA methylation and a set of five histone H3 PTMs in the model hornwort *Anthoceros agrestis* [37] and compared it to *P. patens* and *M. polymorpha*. We conclude that the chromatin organization of bryophytes is distinct from that described in other groups of eukaryotes. It is a patchwork of transposable elements and genes forming small units of euchromatin and heterochromatin without segregation of large heterochromatin domains around centromeres.

## Results

### Chromatin profiling of *Anthoceros agrestis*

We used Enzymatic Methyl-seq [38] to obtain a genome wide profile of 5-methyl cytosines from four week old vegetative tissue after transfer to new growth media of *A. agrestis*. Chromatin immunoprecipitation coupled with DNA sequencing (ChIP-seq) was applied to the same tissue to obtain genomic profiles of five histone PTMs (H3K4me3, H3K36me3, H3K9me1, H3K27me1, and H3K27me3) and H3. Although H3K9me2 is often considered as a mark for constitutive heterochromatin, we did not detect peaks of enrichment for this PTM in *A. agrestis*. Instead, we used H3K9me1 as a mark for constitutive heterochromatin as this PTM shows similar coverage as H3K9me2 in *M. polymorpha* and represents broadly constitutive heterochromatin in *Arabidopsis thaliana*. Peaks from each replicate of all five marks exhibited a significant overlap between each other (Additional File 1: Table S1). Furthermore, biological replicates clustered together in a Pearson correlation matrix (Additional File 2: Fig. S1A) and the profiles of PTMs typically considered repressive (H3K9me1, H3K27me1 and H3K27me3) or active (H3K4me3 and H3K36me3) formed two clusters of high similarity (Additional File 2: Fig. S1A), showing the robustness of our data. In addition, we re-annotated transposable elements (TEs) in the *A. agrestis* Oxford strain and identified 88,959 TEs, including 1155 intact TEs belonging to various TE families (Additional File 3: Supplementary Data 1). A large majority of the annotated TEs were relatively short and, apart from MITEs, were primarily fragments of intact TEs. Peaks of H3K9me1 and H3K27me1 were associated with high levels of DNA methylation in both CG and CHG contexts and primarily associated with various types of TEs but also some protein coding genes (PCGs) (Figs. 1A and B). Peaks of H3K4me3 were enriched in protein coding genes (PCGs) and with very low levels of DNA methylation (Figs. 1A and B). Unlike these associations that are observed in other model land plants, peaks of H3K36me3 and H3K27me3 were not only present on PCGs but also on TEs with modest levels of cytosine methylation (Figs. 1A and B). While in the *A. agrestis* genome many of the associations between genomic features and PTMs were identical to those reported in other model plants, this overview suggested additional associations present in hornworts.

**Figure 1.**
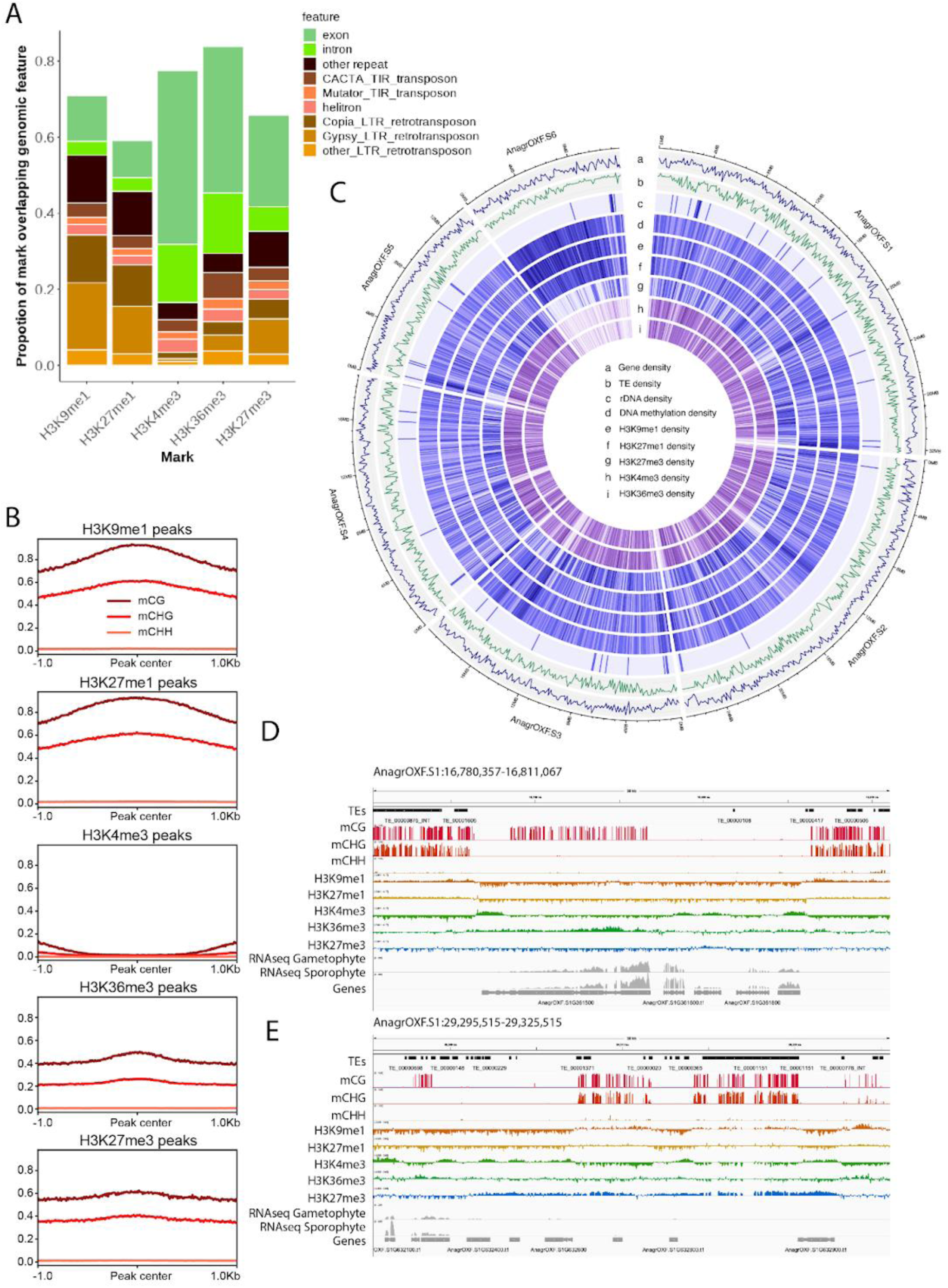
Association of PTMs with genomic features. (A) Distribution of PTMs over genomic features. The total length of PTMs overlapping specified genomic features was divided by the total length of PTM peaks to determine each proportion. Features which cover less than 0.5% of PTM peaks are not shown. (B) Profile plot of CG, CHG, and CHH methylation levels over PTM peaks. Sequences 1 kb upstream and downstream of the peak center are included. Average methylation over 10 bp bins is plotted. (C) Genomic features of the *A. agrestis* genome. The Circos plot illustrates the genomic features of the *A. agrestis* genome using rings to display different information with a window size of 100 kb. The rings represent the following features: a. Gene density b. Transposable element (TE) density c. Ribosomal DNA (rDNA) density d. DNA methylation density e. H3K9me1 peak density f. H3K27me1 peak density g. H3K27me3 peak density h. H3K27me3 peak density i. H3K36me3 peak density. (D) and (E) Integrative Genomics Viewer (IGV) browser screenshot demonstrating vicinity of TEs covered by H3K9me1 and expressed genes (A) and TEs covered by H3K27me3 (B). The regions shown are 30 kb in length from the scaffold AnagrOXF.S1. PTM tracks are bigwig files scaled by H3 coverage in 10-bp windows. DNA methylation tracks are bigwig files showing methylation levels of each cytosine site covered by at least 10 reads. ‘‘TEs’’ and ‘‘Genes’’ tracks are annotation files for TEs and genes, respectively. ‘‘RNA-seq’’ tracks are bigwigs of mapped RNA-seq reads from gametophyte tissue and sporophyte tissue [39]. Scales are noted in square brackets in each track.

To investigate the chromosome level organization of the chromatin landscape in *A. agrestis*, we upgraded the existing genome assembly of the *A. agrestis* Oxford strain [39] by generating additional nanopore long reads as well as Hi-C data. The contig N50 of the new assembly increased from 1.8 to 7.0 Mb. Furthermore, contigs were placed into six large scaffolds corresponding to the six chromosomes in *A. agrestis,* representing 98% of the total assembly with a BUSCO completeness of 92% and LTR assembly index of 19.8. Using this assembly, we plotted densities of protein coding genes (PCGs), TEs, and PTMs to see the distribution of these features over chromosomes (Fig. 1C). Overall, genes and TEs were evenly distributed over the chromosome of *A. agrestis* Oxford strain as reported previously in the *A. agrestis* Bonn strain [39]. Corresponding to this pattern, all PTMs were evenly distributed evenly over the chromosomes, except for chromosome 6 discussed below (Fig. 1C). We observed active PCGs covered by euchromatic marks and surrounded by TEs covered by heterochromatic marks (Fig. 1D). We also observed relatively long genomic regions that included both PCGs and TEs covered by H3K27me3 (Fig. 1E). This even distribution of PTMs, genes, and TEs observed in chromosomes one to five was comparable to the chromatin organization in *M. polymorpha* and *P. patens*.

In contrast to the even distribution of PCGs, TEs and associated PTMs over these five chromosomes, there were fewer PCGs and more TEs on chromosome 6 and most of this this chromosome was covered by DNA methylation, H3K9me1 and H3K27me1 (Fig. 1C, Table1, Additional File 2: Figs. S1B and S1C). In the Hi-C contact map, we observed that chromosome 6 has more intra-chromosomal contacts and fewer inter-chromosomal contacts compared to the other chromosomes (Fig. 2A). We also calculated the densities of each PTM per chromosome and found that chromosome 6 was mostly occupied by constitutive heterochromatin with high densities of DNA methylation, H3K9me1 and H3K27me1 and low densities of H3K4me3, H3K36me3 and H3K27me3 compared to the other chromosomes (Fig. 2B). Corresponding to these heterochromatic characteristics, PCGs on chromosome 6 were expressed at a lower level than PCGs on the other five chromosomes (Fig. 2C). The characteristics of chromosome 6 were similar to those of the sex chromosomes in the dioicous species *M. polymorpha* [32]. However, chromosome 6 did not show an enrichment of orthologs of genes present on Marchantia sex chromosomes (Table 2, Additional File 4: Supplementary Data 2). *Anthoceros agrestis* is monoicous and the association of chromosome 6 with constitutive heterochromatin remains enigmatic. To explore relationships between higher order chromosomal structure and epigenetic marks, we annotated topologically associated domains (TADs) and split the genome into two compartments (active A compartment and inactive B compartment) based on the Hi-C data and calculated the density of each epigenetic mark in TADs or in A/B compartments. We observed a slight enrichment of repressive marks (DNA methylation, H3K9me1, H3K27me1 and H3K27me3) and TEs inside the TADs and a slight enrichment of active marks (H3K4me3 and H3K36me3) and PCGs at TAD boundaries (Fig. 2D and Additional File 2: Fig. S1D). The A compartments showed enrichment of active marks while repressive marks identified the B compartments (Fig. 2E). The centromeres of the chromosomes of *A. agrestis* were not conspicuous based on the Hi-C map and were not marked by a strong accumulation of transposable elements (Figs. 1C and 2A). These features of TAD, A/B compartments and centromeres were similar to those described in *M. polymorpha* [32] and *P. patens* [40]. We conclude that the overall genome organization of bryophytes shows similar features.

**Figure 2.**
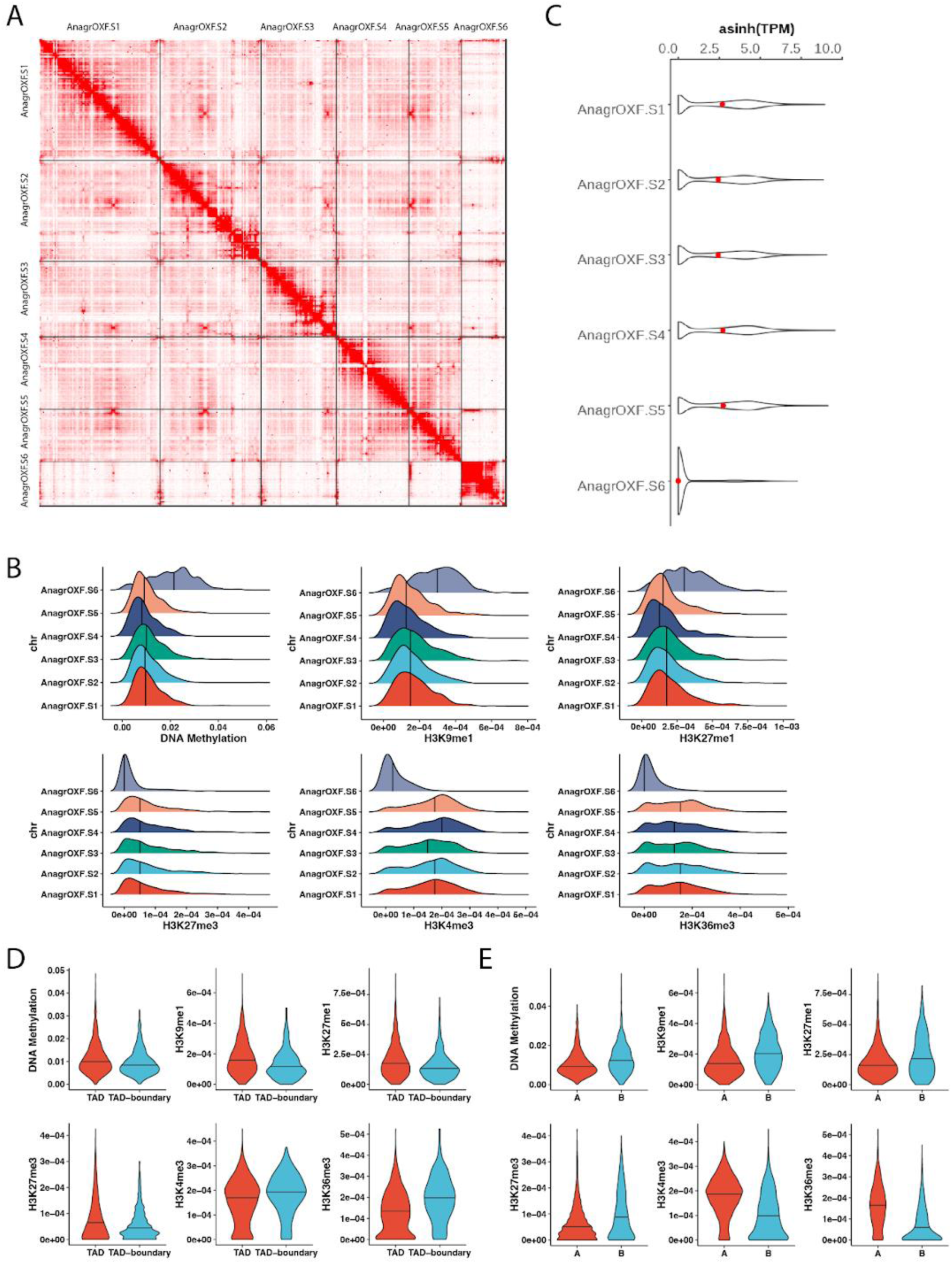
Higher order structure of *A. agrestis* chromosomes. (A) Hi-C contact heatmap of chromosomes. The blocks were utilized to symbolize the signal linked with the two contact positions. The color depth corresponds to the intensity of the interaction between DNA molecules. The darker shades indicate a stronger interaction between them. (B) Density plots showing distributions of epigenetic marks per chromosome. The density of DNA methylation or histone PTMs were calculated as the number of DNA methylation sites or numbers of peaks of histone PTMs in each 40 kb window, divided by window size (40 Kb), and plotted for each chromosome. The median value for each chromosome is represented by a solid vertical line. (C) Violin plot showing expression level of PCGs per chromosome. Width is relative to PCG density. Red dots indicate median expression values. (D) Violin plots showing density distribution of epigenetic marks in TADs and TAD boundaries. The density of epigenetic marks was calculated as the number of histone PTM peaks or DNA methylation sites in each 40 kb window, divided by window length, and plotted for TADs and TAD boundaries. The median value is represented by a solid horizontal line. (E) Violin plots showing density distribution of epigenetic marks in different compartments. The density of epigenetic marks calculated as above is plotted against A or B compartment. The median value is represented by a solid horizontal line.

**Table 1.** Summary of PCG and TE proportions per chromosome.

|  | total<br>length<br>(Mbp) | PCG<br>occupancy<br>(Mbp) | PCG<br>proportion<br>(%) | PCG<br>counts | TE occupancy<br>(Mbp) | TE<br>proportion(%) | TE<br>counts |
| --- | --- | --- | --- | --- | --- | --- | --- |
| AnagrOXF.S1 | 32.3 | 11.8 | 36.5 | 6864 | 12.1 | 37.5 | 21951 |
| AnagrOXF.S2 | 27.0 | 9.8 | 36.5 | 5827 | 10.8 | 39.9 | 18853 |
| AnagrOXF.S3 | 20.0 | 7.0 | 34.9 | 4136 | 8.4 | 42.0 | 14600 |
| AnagrOXF.S4 | 19.2 | 7.9 | 40.9 | 4455 | 6.5 | 33.7 | 11756 |
| AnagrOXF.S5 | 14.1 | 5.6 | 40.0 | 3173 | 4.5 | 32.3 | 8147 |
| AnagrOXF.S6 | 11.8 | 2.5 | 21.6 | 2170 | 8.1 | 68.8 | 10960 |
| total | 124.4 | 44.7 | 35.9 | 26625 | 50.4 | 40.5 | 86267 |

**Table 2.**
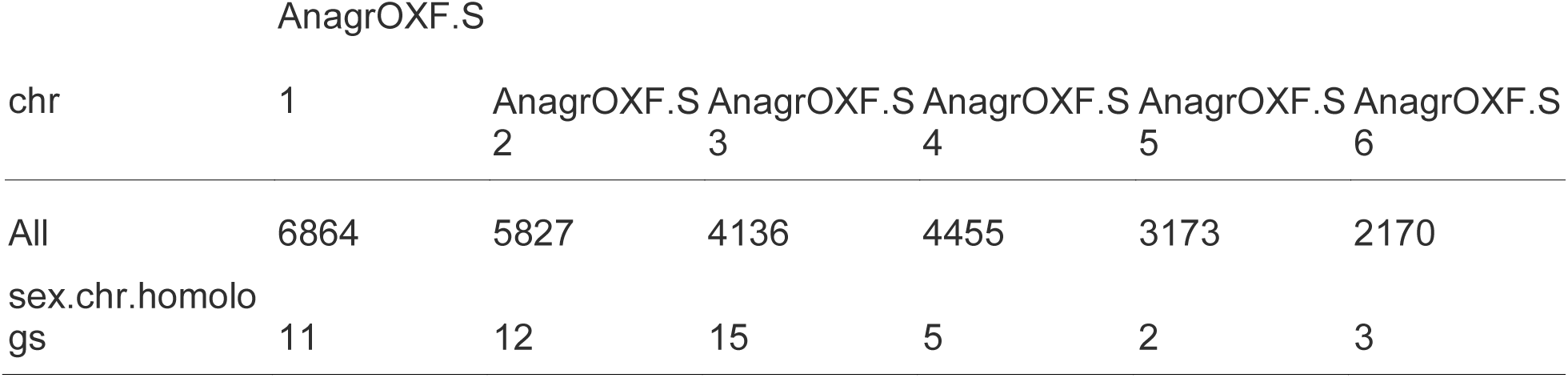
number of homologs of sex chromosome genes in *M. polymorpha* per chromosome.

### Association of histone PTMs and DNA methylation with protein coding genes

We explored preferential associations between PTMs and the transcriptional status of PCGs based on their expression levels in vegetative tissue (thallus) from publicly available data [39]. H3K36me3 and H3K4me3 were strongly associated with expressed PCGs (Fig. 3A), while H3K9me1, H3K27me1, and H3K27me3 were associated with repressed PCGs (Fig. 3A). To discover if there were relationships between chromatin profiles and PCGs in *A. agrestis*, we performed unsupervised k-means clustering of chromatin profiles over all PCGs (n=26,601). By calculating Within-Cluster Sum of Square (Additional File 2: Fig. S2A), we defined five major clusters of PCGs (named cluster P1 to P5) which exhibited different chromatin environments (Fig. 3B, Additional File 5: Supplemental Data 3). Cluster P5, comprising 18.5 % of all PCGs (Fig. 3C), were only weakly enriched for any marks examined, likely due to the difficulty of mapping reads to these predicted PCGs (Additional File 2: Fig. S2C), and we did not consider this cluster further. We calculated the average expression level in the other clusters. Clusters P3 and P4 comprised 16.3%, and 40.1% of all PCGs, respectively, and accounted for expressed PCGs (Fig. 3B - D). These PCGs were longer than repressed PCGs (Fig. 3E). In these two clusters, H3K36me3 was enriched over gene bodies and H3K4me3 was enriched at transcription start sites (TSS), as described in other land plants (Fig. 3B and Additional File 2: Fig. S2E). Enrichment of H3K9me1, H3K27me1 and DNA methylation in the promoter region of PCGs differentiated cluster P3 from cluster P4 (Figs. 3B and 3F). Despite the association of these marks with heterochromatin, there was no difference in expression levels of PCGs in these two clusters (Fig. 3D). In these two clusters, gene bodies showed low levels of CG methylation and no cytosine methylation in non-CG contexts (Fig. 3F), similar to what has been described in *P. patens* [41] and *M. polymorpha* [31,42]. We conclude that P3 and P4 comprise euchromatin and expressed PCGs that are devoid of gene body methylation in bryophytes.

**Figure 3.**
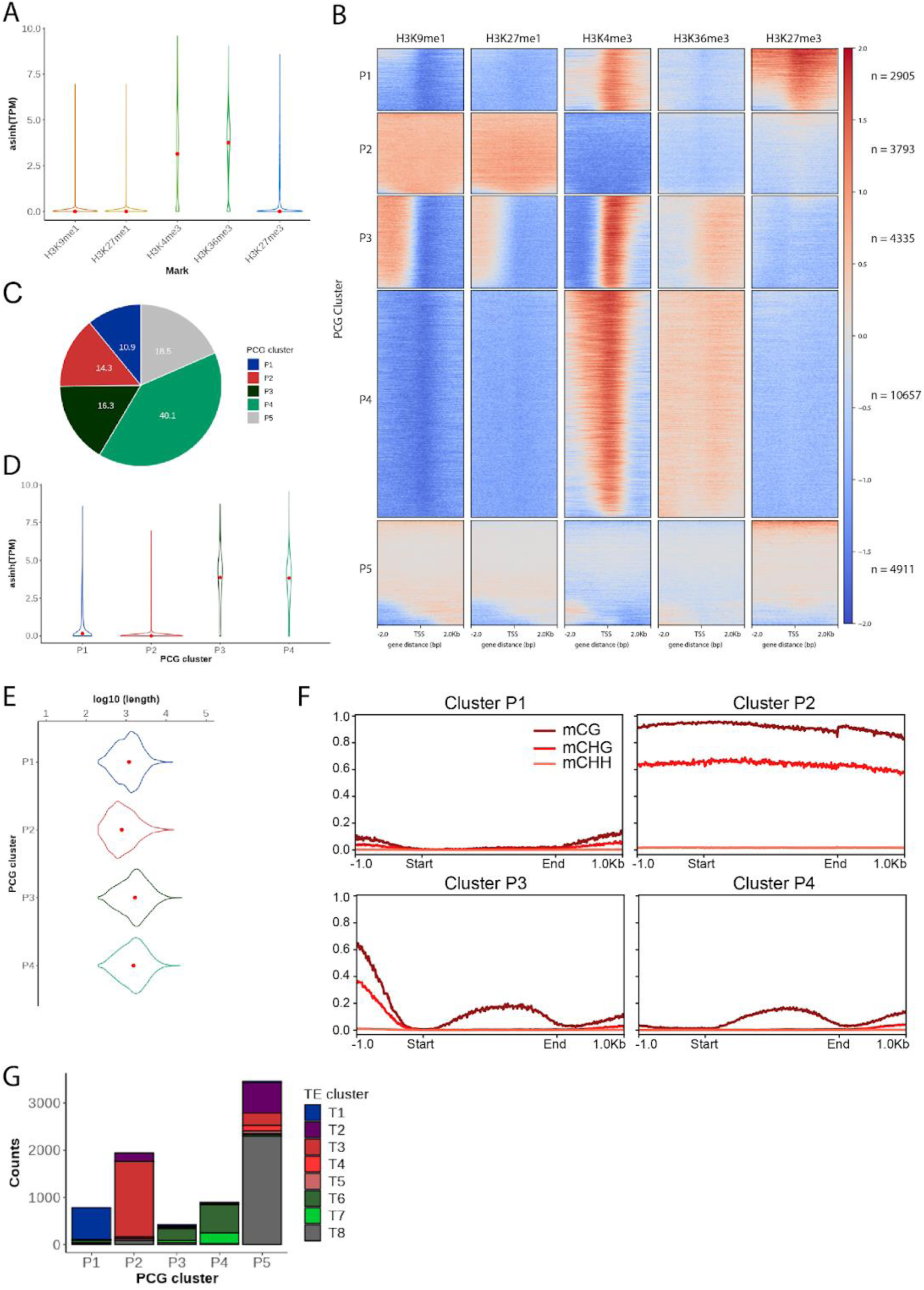
Association of PTMs on PCGs. (A) Violin plot showing expression level of PCGs associated with PTMs. Width is relative to PCG density. Red dots indicate median expression values. (B) Heatmap of k-means clustering of genes based on PTMs. Prevalence of each mark (columns) based on its score normalized against H3 signals per 10 bp bins. Sequences 2 kb upstream and downstream of the start codon are included. Red stands for enrichment and blue for depletion. Each row corresponds to one gene, with multiple genes grouped into blocks that have been defined as clusters P1 through P5. (C) Pie chart showing proportions of PCG clusters in all PCGs.. (D) Violin plot showing expression level of PCGs per PCG cluster. Width is relative to the density of PCGs. Red dots indicate median expression values. (E) Violin plot showing length of PCG per PCG cluster. Width is relative to the density of PCGs. Red dots indicate median values. (F) Profile plot of CG, CHG, and CHH methylation levels over PCGs per PCG cluster. Gene body of each PCG is scaled to 2 kb and sequences 1 kb upstream and downstream are included. Average methylation over 10 bp bins is plotted. (G) Stacked bar chart showing numbers of PCGs overlapped by TEs at least 25% of their length per PCG cluster. Different colors indicate TE clusters defined in Figure 3A.

Non-expressed PCGs formed clusters P1 and P2 (Figs. 3B-D). Cluster P1 contained 10.9% of all PCGs and was characterized by enrichment of H3K27me3 and H3K4me3 and depletion of cytosine methylation in all contexts, and was similar to cluster 1 of *M. polymorpha* PCGs [32]. Cluster P2 contained 14.3% of all PCGs with enrichment of H3K9me1, H3K27me1, and a high level of cytosine methylation in CG and CHG contexts. No comparable chromatin state was observed over PCGs in *M. polymorpha* [32]. PCGs in cluster P2 were shorter than those in other clusters (Fig. 3E). About 60% of PCGs from cluster P2 overlapped with annotated TEs (Fig. 3G), suggesting that these PCGs are coding regions of intact TEs or PCGs that have co-opted coding regions of TEs.

To test if repressed PCGs from clusters P1 and P2 were expressed in non-vegetative tissue, we explored other publicly available transcriptome data sets [39]. While PCGs in clusters P3 and P4 are ubiquitously expressed in both gametophyte and sporophyte, we found that 15% of PCGs in cluster P1 are repressed in gametophytes but expressed in sporophytes, suggesting H3K27me3 functions as a facultative heterochromatic mark to regulate expression of genes regulating the distinct developmental programs of the haploid and diploid phases of the life cycle of *A. agrestis* (Additional File 2: Figs. S3 A, C and D). Surprisingly, 9.3% of PCGs in cluster P2 were repressed in gametophytes but expressed in sporophyte stage, suggesting that H3K9me1 and/or H3K27me1 also behave as facultative heterochromatic marks in *A. agrestis* (Additional File 2: Fig. S3B). These PCGs did not overlap with annotated TEs and encoded proteins such as the GDSL lipase involved in the production of cuticle [43], transporters, and pectin lyase, which altogether might have physiological functions during the sporophyte stage and its adaptation to a terrestrial lifestyle (Additional File 6: Supplementary Data 4).

We conclude that the patterns of enrichment of H3K4me3 and H3K36me3 over expressed genes in euchromatin are broadly conserved in land plants, while in bryophytes facultative heterochromatin associates H3K27me3 with modifications H3K9me1 and H3K27me1 that are usually distinctive of constitutive heterochromatin and TEs. In the lineage of land plants this form of heterochromatin is unique to bryophytes and represses gene expression in a context dependent manner.

### Association of histone PTMs and DNA methylation on TEs

To address the relationships between PTMs and TEs, we performed k-means clustering of histone H3 PTMs over TEs. We defined eight major clusters of TEs showing different chromatin environments (clusters T1 to T8 in Fig. 4A, Additional File 2: Fig. S2B and Additional File 7: Supplemental Data 5). Cluster T8, containing 20.4% of all TEs (Fig. 4B), was weakly enriched for all marks examined, likely due to the difficulty of mapping (Additional File 2: Fig. S2D). Compared to the other clusters, cluster T8 contained longer TEs, which could explain the lower mappability (Fig. 4C). These TEs showed high levels of DNA methylation (Fig. 4D) and half of them were retrotransposons (Fig. 4E). Clusters T3, T4 and T5 contained 23.7%, 6%, and 5.6% of all TEs respectively (Fig. 4B). These TEs were covered with H3K9me1, H3K27me1, and CG and CHG methylation (Figs. 4A and 4D). Cluster T3 contained longer TEs compared to those in clusters T4 and T5 (Fig. 4C) and were more enriched in LTR families (Fig. 4E). Clusters T4 and T5 were enriched for H3K4me3 in the upstream and downstream regions, respectively, and contained TEs shorter than TEs in cluster T3. These two clusters were enriched in DNA transposons (Fig. 4E). Hence, typical heterochromatin occupied TEs from clusters T3, T4, T5, and T8 representing slightly above half of all TEs of *A. agrestis*.

**Figure 4.**
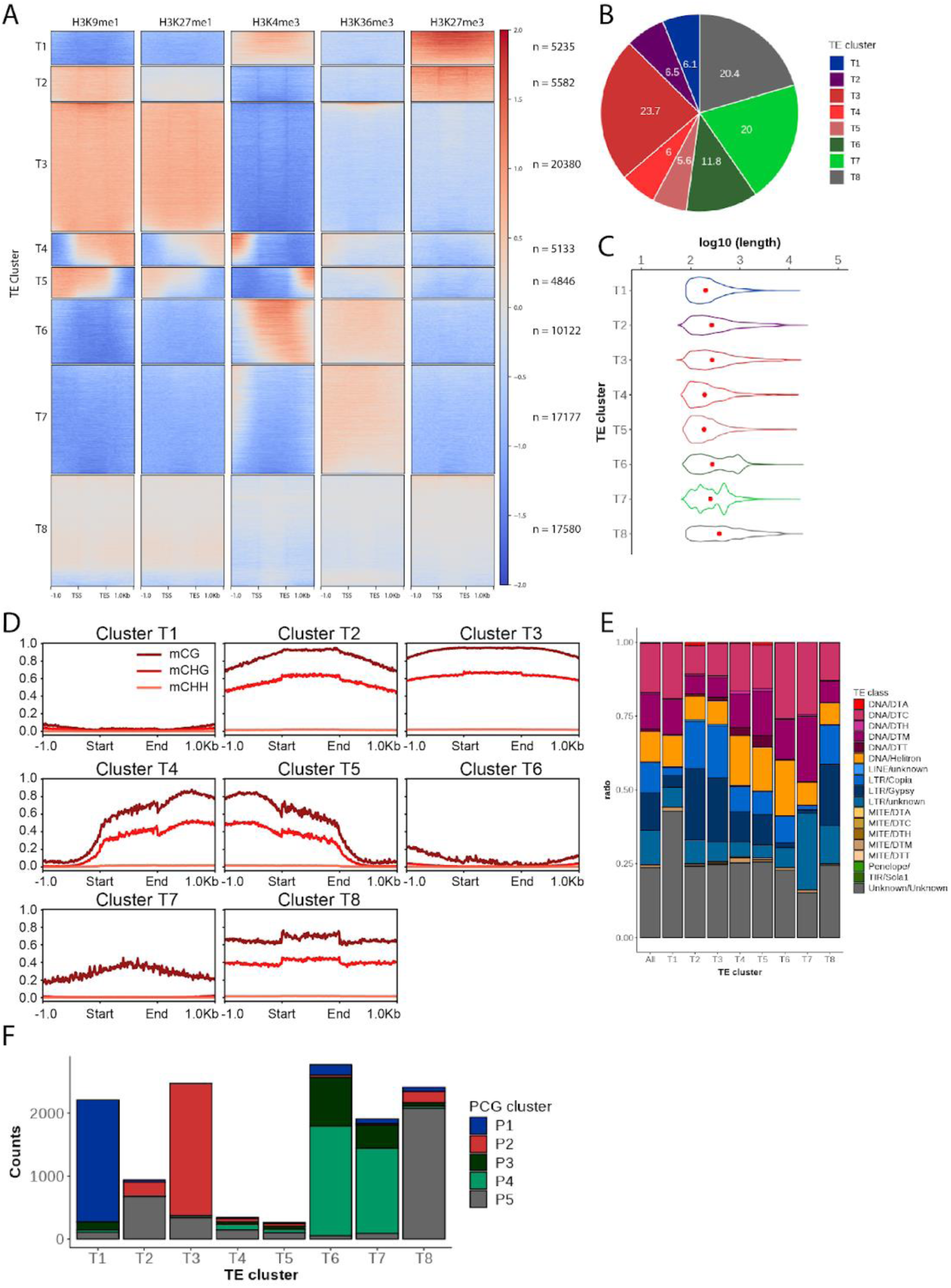
Association of PTMs on TEs. (A) Heatmap of k-means clustering of TEs based on PTMs. Prevalence of each PTM (columns) based on its score normalized against H3 signals per 10 bp bins. Each TE annotation is scaled to 1 kb and sequences 1 kb upstream and downstream are included. Red color stands for enrichment and blue for depletion. Each row corresponds to one TE, with multiple TEs grouped into blocks that have been defined as clusters T1 through T8. (B) Pie chart showing proportions of TE clusters in all TEs. (C) Violin plot showing length of TEs per TE cluster. Width is relative to the density of TEs. Red dots indicate median values. (D) Profile plot of CG, CHG, and CHH methylation levels over TEs per TE cluster. Each TE annotation is scaled to 1 kb and sequences 1 kb upstream and downstream are included. Average methylation over 10 bp bins is plotted. (E) Stacked bar chart indicating proportions of TE families in each TE cluster (T1 to T8) in comparison with TE family proportion in the entire genome (All). (F) Stacked bar chart showing numbers of TEs overlapped by PCGs at least 25% of their length per TE cluster. Different colors indicate PCG clusters defined in Figure 2B.

By contrast, chromatin states distinct from the typical constitutive heterochromatin were observed on TEs from clusters T1, T2, T6, and T7. Clusters T1 and T2 comprised 6.1% and 6.5% of all TEs, respectively (Fig. 4B). TEs from these clusters were associated with H3K27me3 and either H3K4me3 (T1) or H3K9me1 and DNA methylation (T2; Figs. 4A and 4D). Cluster T1 contained shorter TEs and was enriched in unclassified TEs, while cluster T2 contained longer TEs and was enriched in LTR TEs (Figs. 4C and 4E). Clusters T6 and T7 comprised 11.6% and 20% of all TEs and were covered with euchromatic marks (H3K36me3 and H3K4me3) and low levels of CG and CHG methylation (Figs 4A, 4B, and 4D). These TEs were relatively short and enriched amongst DNA transposons (Figs. 4C and 4E). One fourth of T6 and 12% of T7 overlapped with PCGs (Fig. 4F), suggesting that parts of them were located inside PCGs. In conclusion, a relatively small fraction of TEs is covered by marks of constitutive heterochromatin (H3K9me1 and H3K27me1) and is associated with high levels of DNA methylation. These TEs are primarily LTR retrotransposons (Additional File 2: Fig S4). We conclude that only a small fraction of TEs covered with constitutive heterochromatin still have the potential to transpose. The large fraction of TEs associated with facultative heterochromatin or euchromatin might participate in the transcriptional regulation of PCGs in constitutive heterochromatin and in euchromatin.

### Positional relationship between TEs and PCGs

Overall, the genome of *A. agrestis* comprises 20% PCGs and 80% TEs that do not segregate in large domains of constitutive heterochromatin but are rather interspersed with PCGs at the chromosome scale. It is thus possible that PCGs or TEs form small clusters. To answer this question we first identified the probability of a PCG or a TE belonging to a specific cluster to be surrounded by another PCG or TE upstream or downstream (Figs. 5A and 5B). We removed overlapping features between the PCG annotation and TE annotation and used only non-overlapping annotations to call the nature (PCG or TE) of the closest neighbors of each PCG and TE per cluster and plotted PCG:TE ratio. This ratio was compared to the overall ratio of PCG/TE (1/4). We observed that PCGs tended to cluster together (Fig. 5A). This tendency was the strongest in PCGs of cluster P4 that constituted small islands of euchromatin (Additional File 2: Fig. S5A) and the weakest in PCGs of cluster P2 covered with constitutive heterochromatin that were surrounded by TEs (Additional File 2: Fig. S5B). PCGs covered with facultative heterochromatin formed smaller PCGs clusters than euchromatic PCGs. To test if PCGs and surrounding TEs are more likely to share the same type of chromatin environment we established the nature of the chromatin environment of the closest neighbors for each PCG cluster (Figs. 5 C and D) or TE cluster (Figs. 5 E and F). This analysis showed that PCGs (P2) and TEs (T3) covered with constitutive heterochromatin tended to be surrounded by PCGs or TEs also covered with constitutive heterochromatin (Figs. 5 C-F). Similarly, PCGs and TEs in euchromatin (P4, T6, and T7) are surrounded by larger regions of euchromatin. This was also the case for TEs and PCGs covered with facultative heterochromatin (P1, T1, and T2) although these could be surrounded by euchromatic PCGs or TEs with constitutive heterochromatin. By contrast, the region upstream of PCGs from cluster P3 was primarily occupied by TEs from clusters T4 and T5 enriched with H3K9me1 and H3K27me1 (Fig. 5D and Additional File 2: Fig. S5C). Higher PCG ratio were found upstream of TEs from cluster T4 and downstream of TEs from cluster T5. These PCGs were covered by H3K4me3 and H3K36me3 (Fig. 5E). Measuring the distance from each PCG to the closest TE per cluster (Additional File 2: Fig. S5D) and vice versa (Additional File 2: Fig. S5E) revealed clusters of PCGs and TEs. These clusters covered by H3K27me3 were larger than clusters marked by euchromatin and constitutive heterochromatin. We concluded that the genome of *A. agrestis* comprises clusters of TEs and PCGs sharing the same chromatin environment, except for P3 euchromatic PCGs with an upstream region enriched in TEs and constitutive heterochromatin.

**Figure 5.**
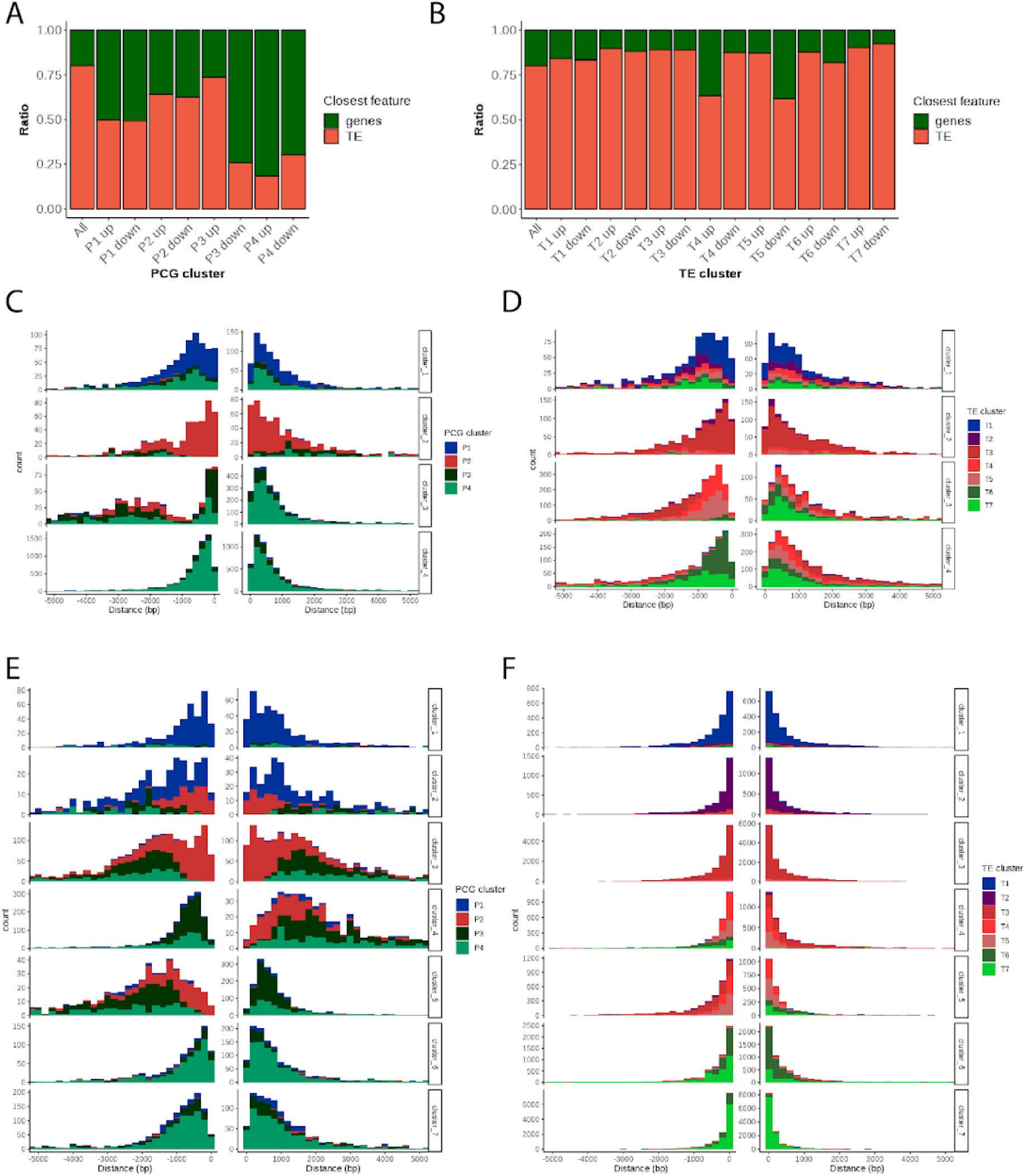
Positional relationships between PCGs and TEs. (A) and (B) Stacked bar chart showing the proportion of closest genomic features (PCGs or TEs) upstream or downstream of PCGs (P1 to P4 in A) and TEs (T1 to T8 in B) per cluster in comparison with the proportion in the entire genome (All). (C), (D), (E) and (F) Histograms showing distance from PCGs (C and D) or TEs (E and F) to the closest PCGs (C and E) or TEs (D and F) per cluster.

**Figure 6.**
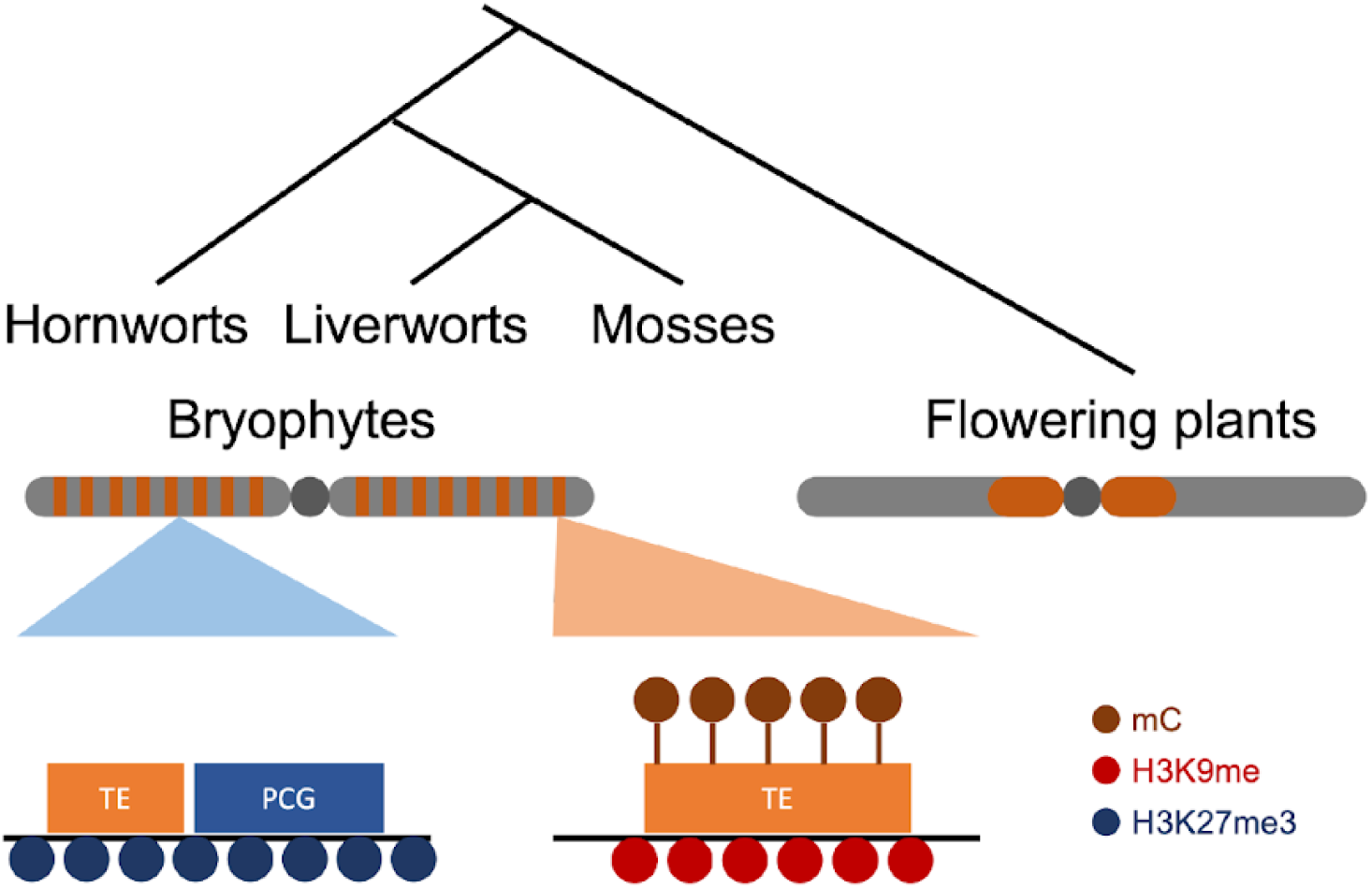
Chromatin organization in the common ancestor of bryophytes. Ancestral chromatin organization in bryophytes is inferred by chromatin synapomorphies shared by three bryophyte species. In contrast to the large pericentromeric heterochromatin observed in flowering plants, the constitutive heterochromatin of bryophytes forms small units and is scattered over chromosomes in bryophytes. These constitutive heterochromatin marked by H3K9me and DNA methylation primarily consist of TEs. In addition, facultative heterochromatin marked by H3K27me3 contains not only PCGs but also TEs.

## Discussion

In flowering plants, TEs are enriched in pericentromeric heterochromatin while PCGs are enriched along chromosomal arms [6,13]. By contrast, both TEs and PCGs are evenly distributed over the entire chromosomes in bryophytes [32,39,44,45]. Consistently euchromatin, facultative and constitutive heterochromatin are also evenly distributed over the chromosomes of *M. polymorpha* and *P. patens* [30,32]. In this study, we demonstrated that an even distribution of the chromatin modifications shapes the global architecture of the genome of the hornwort *A. agrestis* similar to *M. polymorpha* and *P. patens* [30,32]. The conservation of genome size and number of chromosomes amongst hornworts (generally 6;[46] and liverworts (always 9 ;[47] also supports a general conservation of genome architecture across the bryophytes, but there might be deviations to this rule in mosses with much larger genomes.

H3K9 methylation forms constitutive heterochromatin that represses the expression of TEs in many eukaryotic species [23]. In addition, this mark strongly co-occurs with H3K27me1 and DNA methylation in angiosperms [26,27]. In all model bryophytes, the majority of TEs are also associated with H3K9me1 and H3K27me1, in agreement with the presence of SUVH4/5/6 and ATXR5/6 orthologs in bryophytes. In *M. polymorpha* the ortholog of *A. thaliana* MET1 maintains CG methylation and is involved in silencing TEs [31]. In *P. patens*, CHG methylation is maintained by an ortholog of *A. thaliana* CHROMOMETHYLASE 3 [48] and CHH methylation is primarily deposited by the *de novo* methyltransferase DNMT3. *P. patens* lacks orthologs of the methyltransferases DRM1, DRM2 [29] or CMT2, which are collectively responsible for CHH deposition and contribute to TE silencing in *A. thaliana* [48,49]. We identified orthologs of MET1 and CMT in *A. agrestis*, in agreement with the presence of CG and CHG methylation (Table 3). However, although DRM orthologs are present in *A. agrestis*, the near complete absence of CHH methylation suggests that the DRM ortholog is inactive in vegetative haploid tissues. DNA methylation in CHG and CG is primarily observed in TEs, suggesting the absence of gene body methylation on PCGs as described in *A. thaliana.* Such reduction or absence of gene body methylation was also reported in *M. polymorpha* [33] and *P. patens* [48] and is thus a common feature in bryophytes.

**Table 3.**
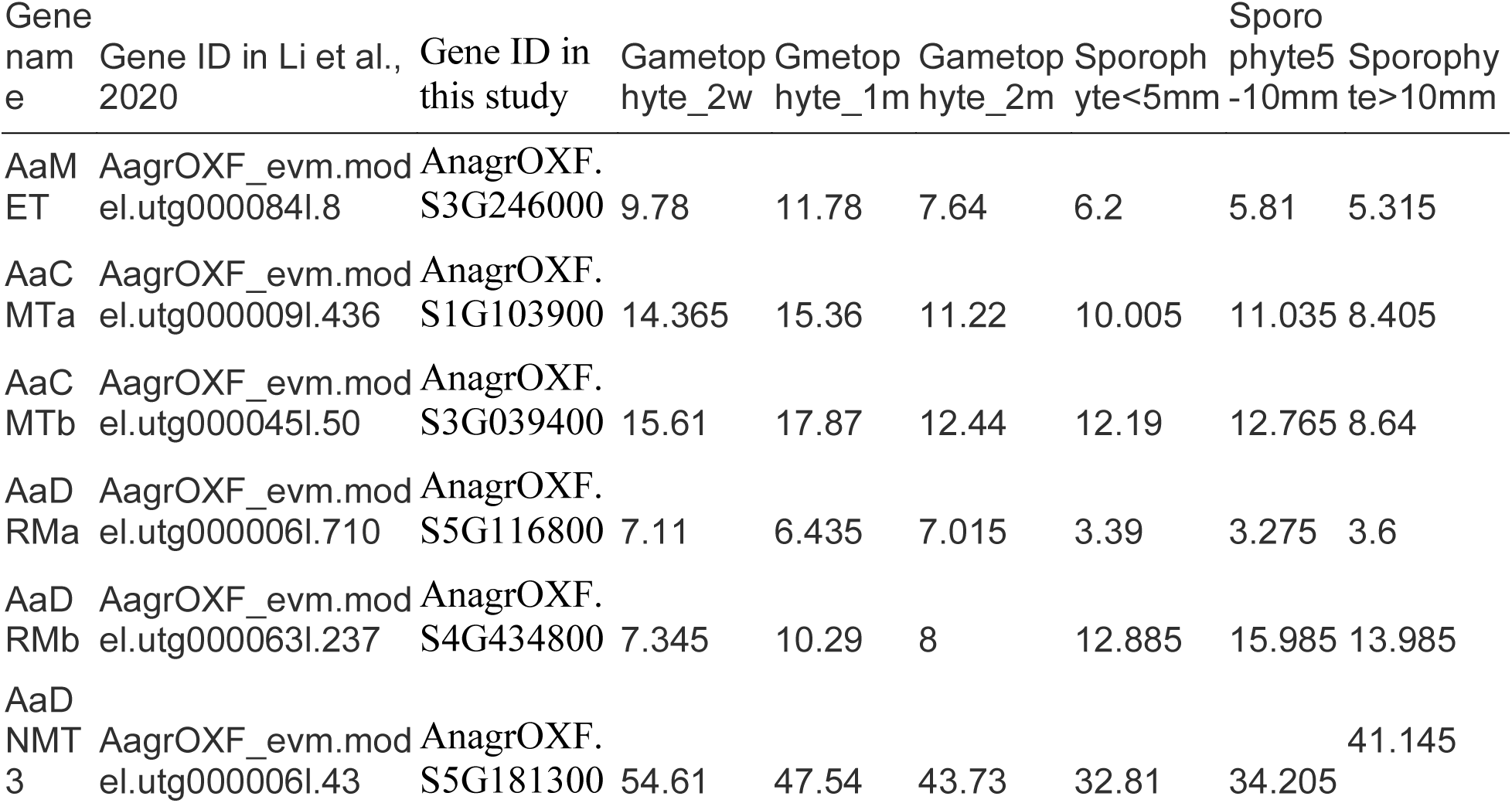
Expression levels of homologs of DNA methyltransferase in *A. agrestis*. Gene name, gene ID and averaged TPM value in each developmental stage are shown.

Despite the overall conservation of chromatin environments in bryophytes, several features are specific to *A. agrestis*. First, a significant fraction of PCGs are surrounded by TEs forming small clusters covered by constitutive heterochromatin (H3K9me1 and H3K27me1). Strikingly, half of all TEs show very low levels of DNA methylation, suggesting that TEs play an important role outside of constitutive heterochromatin in *A. agrestis*. Large clusters of facultative heterochromatin comprise TEs and PCGs covered by H3K27me3. This is reminiscent of the association between TEs and H3K27me3 in *M. polymorpha*, further supporting a shared role of H3K27me3 in silencing TEs across a broad range of eukaryotes [50]. On the other hand, short TE fragments covered by euchromatic marks (H3K4me3 and H3K36me3) are located near or inside PCGs. This strong association between TEs and euchromatin is not observed in *M. polymorpha* or *A. thaliana* and might result from the reduced size of the *A. agrestis* genome (130 Mb and 26,000 PCGs) compared to *M. polymorpha* (230Mb and 20,000 PCGs).

Our results also suggest that facultative heterochromatin of hornworts is not only marked by H3K27me3 but also by H3K9me1 together with H3K27me1 on PCGs that are specifically expressed in sporophytes. Almost half of all TEs in *A. agrestis* are present in euchromatin or facultative heterochromatin, indicating a large degree of co-option of TEs to PCGs and/or their regulatory sequences. Such co-option has been reported mammals [51] and in the pollen of *A. thaliana*, where reprogramming of H3K9me2 and DNA methylation is associated loci in vegetative nuclei promote expression of genes required for pollen tube growth [52,53]. The scattered location of constitutive heterochromatin in bryophytes contrasts with the position of heterochromatin around point centromeres in flowering plants with small genomes [54,55]. It has been proposed that heterochromatin participates in the recruitment of centromeric proteins in fission yeast, drosophila, and *A. thaliana* [56,57]. By contrast, bryophytes might have evolved mechanisms of centromere recruitment independent of constitutive heterochromatin and distinct from that of flowering plants. This mechanism could be related to the pervasive co-option of TEs in bryophytes that selected their recruitment outside of centromeres but not in discrete domains surrounding centromeres. The recent chromosome-scale assembly of the genome of the streptophytic alga *Zygnema cf cylindricum* [58] shows an organization of TEs and PCGs similar to that of bryophytes, suggesting that the chromatin organization described for bryophytes is representative of the common ancestor of land plants.

## Conclusions

We conclude that chromatin synapomorphies shared by *A. agrestis* and either *M. polymorpha* or *P. patens* represent ancestral features of chromatin in bryophytes: an absence of segregation of large domains of heterochromatin to the exception of constitutive heterochromatin of the sex chromosome, small clusters of PCGs and TEs sharing the same chromatin state and forming TADs, an absence of DNA methylation on gene bodies and a separation between constitutive and facultative heterochromatin less demarcated than in flowering plants.

## Methods

### Plant materials

*Anthoceros agrestis* strain Oxford was cultured on 0.5 Gamborg’s B5 medium solidified with 1% agar under continuous white light at 22°C.

### Chromosomal assembly of *A. agrestis* Oxford strain genome

DNA extracted from axenic cultures was sequenced using Oxford Nanopore MinION (ONT) and a total of 11.8 Gb of long-read sequences were obtained. A draft assembly was constructed from the longest ONT reads that yield 50X coverage using Flye 2.9 [59]. The assembly was error-corrected with Pilon 1.24 [60] using Illumina genomic data from [39]. Hi-C data were generated and used to scaffold the assembly by Phase Genomics (Seattle, WA, USA). Gaps between contigs in the scaffolded assembly were filled by TGS-GapCloser [61] using ONT DNA sequences. One 7.4 Mb contig with an unusually high GC content (72%), very low coverage in the Illumina dataset (<1X), and high identity BLAST hits to Actinobacteria was removed as a likely contaminant. A custom repetitive element library was constructed with EDTA 2.0 [62] and used to mask repeats throughout the genome. BRAKER2 v2.1.5 [63] was used with a combination of RNAseq data from [39] mapped to the repeat-masked genome with HISAT2 [64], and genes previously predicted in hornworts [39,45]. Genome and annotation completeness were assessed by BUSCO v5 with viridiplantae_odb10 dataset [65], and LTR assembly index [66].

### Annotation of TEs in *A. agrestis*

Transposable elements of *A. agrestis* were annotated using EDTA 1.9.9 [62], which incorporates several tools, including LTRharvest, LTR_FINDER, LTR_retriever, Generic Repeat Finder, TIR-Learner, MITE-Hunter, HelitronScanner, and RepeatMasker. All tools were adjusted to EDTA with proper filters and parameters. Final, non-redundant TE libraries were produced by removing nested insertions and protein-coding genes with EDTA using customized scripts.

### Chromatin profiling of *A. agrestis* using ChIP-seq

ChIP experiments were performed using a previously described protocol with some modifications [67]. Four weeks old gametophyte tissue of *A. agrestis* were collected and crosslinked using 1% paraformaldehyde in 1x PBS under vacuum on ice for 10 minutes. The cross-linking reaction was quenched by adding 2 M glycine under vacuum on ice for 10 minutes. Excess solution was removed from cross-linked tissue by blotting with paper towels. Cross-linked tissue was then snap frozen in liquid nitrogen and ground to a fine powder using mortar and pestle. The powder was transferred into a 50 ml plastic tube and suspended in 40 ml of MP1 buffer (10 mM MES-KOH buffer pH 5.3, 10 mM NaCl, 10 mM KCl, 0.4 M sucrose, 2% (w/v) PVP-10, 10 mM MgCl2, 10 mM 2-mercaptoethanol, 6 mM EGTA, 1× cOmplete protease inhibitor cocktail). Suspended samples were then filtered twice through one layer of Miracloth, once through a 40 μm nylon mesh once, and twice through a10 μm nylon mesh. Filtered samples were centrifuged at 3000xg at 4°C for 10 min and the supernatant was discarded. The pellet was washed using 15 ml of MP2 buffer (10 mM MES-KOH buffer pH 5.3, 10 mM NaCl, 10 mM KCl, 0.25 M sucrose, 10 mM MgCl2, 10 mM 2-mercaptoethanol, 0.2% Triton-X 100, 1× cOmplete protease inhibitor cocktail) three times. The final pellet was then resuspended in 5 ml of MP3 buffer (10 mM MES-KOH buffer pH 5.3, 10 mM NaCl, 10 mM KCl, 1.7 M sucrose, 2 mM MgCl2, 10 mM 2-mercaptoethanol, 1× cOmplete protease inhibitor cocktail) and centrifuged at 16000xg at 4°C for 1 h. After centrifugation, the supernatant was discarded and the pellet was resuspended in 900 μl of covaris buffer (0.1% SDS, 1 mM EDTA, 10 mM Tris-HCl pH 8.0, 1× cOmplete protease inhibitor cocktail). The resuspended pellet containing the chromatin fraction was fragmented using a Covaris E220 High Performance Focused Ultrasonicator for 15 min at 4 °C (duty factor, 5.0; peak incident power, 140.0; cycles per burst, 200) in a 1 ml Covaris milliTUBE. Sheared chromatin was centrifuged at 15000 rpm at 4°C for 10 min and the supernatant was transferred into a new 5 ml tube and diluted by adding 2.7 ml of ChIP dilution buffer. Diluted chromatin was cleaned by incubating with proteinA/G beads (Thermo Fisher Scientific) at 20 rpm spinning at 4°C for 1 h. Beads were removed by magnetic racks and precleared chromatin was separated into 6 tubes and incubated with 1 µg of specific antibodies for histone modifications (listed in Additional File 8: Table S2) at 20 rpm spinning at 4°C overnight. Chromatin bound by antibodies was collected by incubating with protein A/G beads for 3 h. The beads were collected by magnetic racks and washed twice with a low salt wash buffer (20 mM Tris-HCl pH 8.0, 150 mM NaCl, 2 mM EDTA, 1% Triton X-100 and 0.1% SDS), once with a high salt wash buffer (20 mM Tris-HCl pH 8.0, 500 mM NaCl, 2 mM EDTA, 1% Triton X-100 and 0.1% SDS), once with a LiCl wash buffer (10 mM Tris-HCl pH 8.0, 1 mM EDTA, 0.25 M LiCl, 1% IGEPAL CA-630 and 0.1% sodium deoxycholate), and twice with a TE buffer (10 mM Tris-HCl pH 8.0 and 1 mM EDTA). Immunoprecipitated DNA was eluted using 500 µl elution buffer (1% SDS and 0.1 M NaHCO3) at 65°C for 15 min. To reverse cross-link, eluted DNA was mixed with 51 µl of reverse cross-link buffer (40 mM Tris-HCl pH 8.0, 0.2 M NaCl, 10 mM EDTA, 0.04 mg ml−1 proteinase K (Thermo Fisher Scientific)), and incubated at 45°C for 3 h and then at 65°C for 16 h. After cross-link reversal, DNA was treated with 10 µg of RNase A (Thermo Fisher Scientific), incubated at room temperature for 30 min and purified using the MinElute PCR purification kit (Qiagen). ChIP-seq library was generated from ChIPed DNAs using Ovation® Ultralow Library Systems V2 (TECAN). The ChIP-seq libraries were sequenced on illumina Hiseq v4 to generate 50 bp single end reads.

### ChIP-seq data analyses

The bam files of ChIP-seq reads were sorted with SAMtools v1.9 [68] and converted to fastq format using the bamtofastq function of BEDTools v2.27.1 [69], trimmed with Cutadapt v1.18 [70] and aligned to the *A. agrestis* Oxford strain genome assembled in this study using Bowtie2 v2.3.4.2 [71]. Resulting bam files were sorted and indexed with SAMtools v1.9. Reads with MAPQ less than ten were removed with Samtools v1.9 and duplicates were removed with Picard v2.18.27 (http://broadinstitute.github.io/picard/). Pearson correlation matrices were generated using multiBamSummary and plotCorrelation functions in deepTools v3.3.1 [72]. Deduplicated reads from 2 biological replicates were merged. The read coverage of each chromatin mark was normalized against the read coverage of H3 in 10 bp windows with the bamCompare function in deepTools v3.3.1, generating bigwig files. Broad peaks of each chromatin mark were called by using macs2 v2.2.5 with default settings [73]. Overlaps between genomic features and each chromatin mark were calculated by using the intersect function of BEDTools v 2.27.1. The ratio of each genomic feature was calculated by dividing the total length of overlaps by the total length of each chromatin mark.

### Clustering analysis of ChIPseq data

K-means clustering of chromatin marks was performed using deepTools v3.3.1. Matrices were computed using the computeMatrix function of deepTools v3.3.1 with the reference-point sub-command or scale-regions sub-command for PCGs or TEs, respectively, using bigwig files as the input. These matrices were imported into R using profileplyr package and within groups sum of squares were calculated and plotted against number of clusters to estimate optimal numbers of clusters. Heatmaps of matrices were plotted with plotHeatmap with k-means clustering (k = 5 for PCGs and k = 8 for TEs). Cluster assignments can be found in Data S3. Overlaps between PCG annotations and TE annotations were calculated using the intersect function of BEDTools v2.27.1. PCGs are considered as overlapped by TEs when more than 50% of the regions of each PCG are overlapped by each TE and vice versa. Numbers of PCGs overlapped by TEs and TEs overlapped by PCGs per cluster were plotted using the ggplot2 package in R [74]. The distance between the closest TE and PCG pair per cluster was calculated using the closest function of BEDTools.

Genome mappability of the *A. agrestis* Oxford strain genome assembled in this study was calculated by using GenMap [75] with options -K 50 and -E 0. The output bedgraph file was converted to a bigwig file by using bedGraphToBigWig v385 [76]. Average mappability of each PCG and TE was calculated using bigWigAverageOverBed [76] and plotted by using ggplot2 package in R.

To assign the closest features of each PCG or TE, all PCGs and TEs overlapped by TEs and PCGs respectively, were removed by using the intersect function of BEDTools v 2.27.1. Then the nearest genomic feature to each PCG was assigned by comparing the PCG annotation with both PCG and TE annotations using closest function of BEDTools v 2.27.1 with options -io, -mdb all, -D a, -t first and either -id or -iu. The nearest genomic feature to each TE was calculated similarly. These data were plotted by using ggplot2 package in R.

### Gene Expression analysis of *A. agrestis*

Gene expression data from [39] were downloaded from the SRA (PRJNA574453) or ENA (PRJEB34743) and processed with RSEM v1.2.31 [77] and STAR v2.5.2a [78]. Transcript Per Million (TPM) values were averaged from three biological replicates from each condition and used for further analyses. The association of each peak over PCGs was calculated using the intersect function of BEDtools v2.27.1. PCGs were considered as overlapped by each chromatin mark when more than 50% of the regions of each PCG were overlapped by each peak. Average expression levels per chromatin peak were plotted using ggplot2 package in R. Heatmaps of expression levels of PCGs in cluster 1 to 4 over various ages of gametophyte and sporophyte tissues were plotted using pheatmap function in R.

### Genome wide profiling of 5mC in *A. agrestis*

Genomic DNA was extracted from 100 mg of 4 weeks old gametophyte tissue of *A. agrestis* using Nucleon PhytoPure (cytiva). Sequencing libraries for genome wide DNA methylation profiles were generated from 200 ng of genomic DNA using NEBNext® Enzymatic Methyl-seq Kit (New England Biolabs). These libraries were sequenced on an Illumina NextSeq 2000 to generate 100 bp paired end reads.

### EMseq data analysis

The bam files of EM-seq reads were sorted with SAMtools v1.9 and converted to fastq format using bamtofastq function of BEDTools v2.27.1, then trimmed with Trim Galore (https://github.com/FelixKrueger/TrimGalore). A bisulfite converted reference genome was prepared from *A. agrestis* Oxford strain genome sequence using Bismark v0.22.2 [79]. Trimmed reads were mapped to the bisulfite genome using the Bowtie2 option of Bismark v0.22.2. Duplicates were removed using the deduplicate function in Bismark v0.22.2. Cytosine methylation reports were created from deduplicated reads using the bismark_methylation_extractor function in Bismark v0.22.2. Each cytosine which is covered by at least ten reads was used for further analyses. The methylation ratio of each cytosine was calculated and summarized to a bed file. These bed files were converted to bigwig files using bedGraphToBigWig [76] and used as inputs for the computMatrix function in deepTools v3.3.1. Aggregate profile plots of matrices were plotted with the plotProfile function in deepTools v3.3.1.

### Hi-C contact map construction

Hi-C reads were mapped to the *A. agrestis* genome with Juicer by default parameters and visualized using Juicebox [80]. Finally, we obtained DNA contact signal of six pseudo-chromosomes. This visualization provides insight into the spatial organization of the genome by showing the frequency and intensity of physical interactions between different regions.

### Identification of TAD and A/B compartment

HiCExplorer V3.3 [81] was used for the identification of A/B compartment and TAD boundaries. Clean data were initially mapped to the *A. agrestis* genome using bwa mem with parameters ‘-E50 -L0’. Subsequently, dangling end reads, same fragment reads, self-circles reads, and self-ligation reads were removed. Raw Hi-C matrices were generated at resolutions of 20 kb, 40 kb, 100 kb, and 200 kb in h5 format using hicBuildMatrix. To diagnose and correct matrix resolution, the KR method was applied with hicCorrectMatrix. TADs and TAD boundaries were then identified using hicFindTADs, with TAD separation scores calculated at 40 kb resolution using default parameters. Finally, compartmentalization was performed using hicPCA, with compartment A/B assignment indicated by PC1 values from the analysis on correlation maps with the ‘lieberman’ method. Positive and negative values of the first principle components were plotted to indicate high gene density (compartment A) and low gene density (compartment B), respectively.

### Density profile of different genomic features

The density profile was calculated using BEDtools v2.27 [69] as the total number of peaks or DNA methylation sites in each window, divided by window length (40 Kb). Figures were generated using ggplot2.

## Declarations

### Ethics approval and consent to participate

Not applicable.

### Consent for publication

Not applicable.

### Availability of data and materials

Sequencing data generated in this study is available through NCBI Gene Expression Omnibus under the accession number GSE218880. All code used in this study is available upon request.

### Author contributions

T.H. and F.B. conceived and designed the experiments. T.H. performed ChIP-seq and EM-seq in *A. agrestis* with help from S.A.. S.W. annotated TEs in the genomes of *A. agrestis.* T.H. and S.W. performed the analysis of the data with help from E.A. who also performed statistical analyses and curated data. L.D. provided funding of S.W. and F.B. supervised the study. P.S. and F-W.L. provided the genome assembly and gene annotations, HiC data, and its analyses. T.H. and F.B. wrote the manuscript draft with input from all authors.

## Acknowledgements

We thank Sean A. Montgomery for suggestions and critical reading of the manuscript. F.B. acknowledges support from the PlantS and Next Generation Sequencing at the Vienna BioCenter Core Facilities (VBCF), and the Molecular Biology Services.

---

**Figure S1. Quality control of ChIP-seq**

(A) Pearson correlation matrix showing that biological replicates of each mark cluster together.

(B) and (C) Bar plot showing the proportion of PCGs (A) or TEs (B) in each chromosome. Proportions were calculated as the total length of the features in each chromosome is divided by the length of each chromosome.

(D) Violin plots showing density distribution of genomic features (PCGs or TEs) in TADs and TAD boundaries. The density of genomic features was calculated as the number of features in each 40 kb window, divided by window length, and plotted for TADs and TAD boundaries. The median value is represented by a solid horizontal line.

**Figure S2. K-means clustering of ChIP-seq data over PCGs and TEs**

(A) and (B) Within groups sum of squares calculated using the output files of computeMatrix command over PCGs (A or TEs (B) are plotted against numbers of clusters.

(C) and (D) boxplots indicating the mappability of PCGs (C) and TEs (C) per cluster.

(E) Profile plot showing log2 ChIP/H3 enrichment of H3K4me3 and H3K36me3 over PCGs in clusters P3 and P4. Sequences 2 kb upstream and downstream of the start codon are included.

**Figure S3. Expression level of PCGs in gametophyte and sporophyte**

(A), (B), (C) and (D) Heatmaps showing expression levels of PCGs in gametophyte and sporophyte tissue per cluster. Expression levels are indicated by arcsine TPM.

**Figure S4. DNA methylation levels over each TE family**

Profile plot of CG, CHG, and CHH methylation levels over TEs per TE family. Each TE annotation is scaled to 1 kb and sequences 1 kb upstream and downstream are included. Average methylation over 10 bp bins is plotted.

**Figure S5. Distances between PCGs and TEs per cluster**

(A), (B) and (C) Integrative Genomics Viewer (IGV) browser screenshot demonstrating PCGs in the cluster P4 forming a small euchromatic island (A), PCGs in the cluster P2 surrounded by TEs (B) and TEs in the promoter region of PCGs in the cluster P3. The regions shown are 19 kb in length in (A) and (C) or 9.5 kb in (B) from the scaffold AnagrOXF.S1. PTM tracks are bigwig files scaled by H3 coverage in 10-bp windows. DNA methylation tracks are bigwig files showing methylation levels of each cytosine site covered by at least 10 reads. ‘‘TEs’’ and ‘‘Genes’’ tracks are annotation files for TEs and genes, respectively. ‘‘RNA-seq’’ tracks are bigwigs of mapped RNA-seq reads from gametophyte tissue and sporophyte tissue [39]. Scales are noted in square brackets in each track.

(D) Boxplots of distances between each gene and the nearest TE per TE cluster. Briefly, each gene is compared to all TEs belonging to a TE cluster to find its nearest neighbor. Genes are divided based on the gene cluster they belong to. Distances are in kilobase pairs (kbp). Colored boxes represent interquartile range, and lines represent median values. Outliers are not shown.

(E) Boxplot of distances between each TEs and the nearest PCG per PCG cluster. Briefly, each TE is compared to all PCGs belonging to a PCG cluster to find its nearest neighbor. TEs are divided based on the TE cluster they belong to. Distances are in kilobase pairs (kbp). Colored boxes represent interquartile range, and lines represent median values. Outliers are not shown.

