## Supplemental Figures and legends for "The ancestral chromatin landscape of land plants"

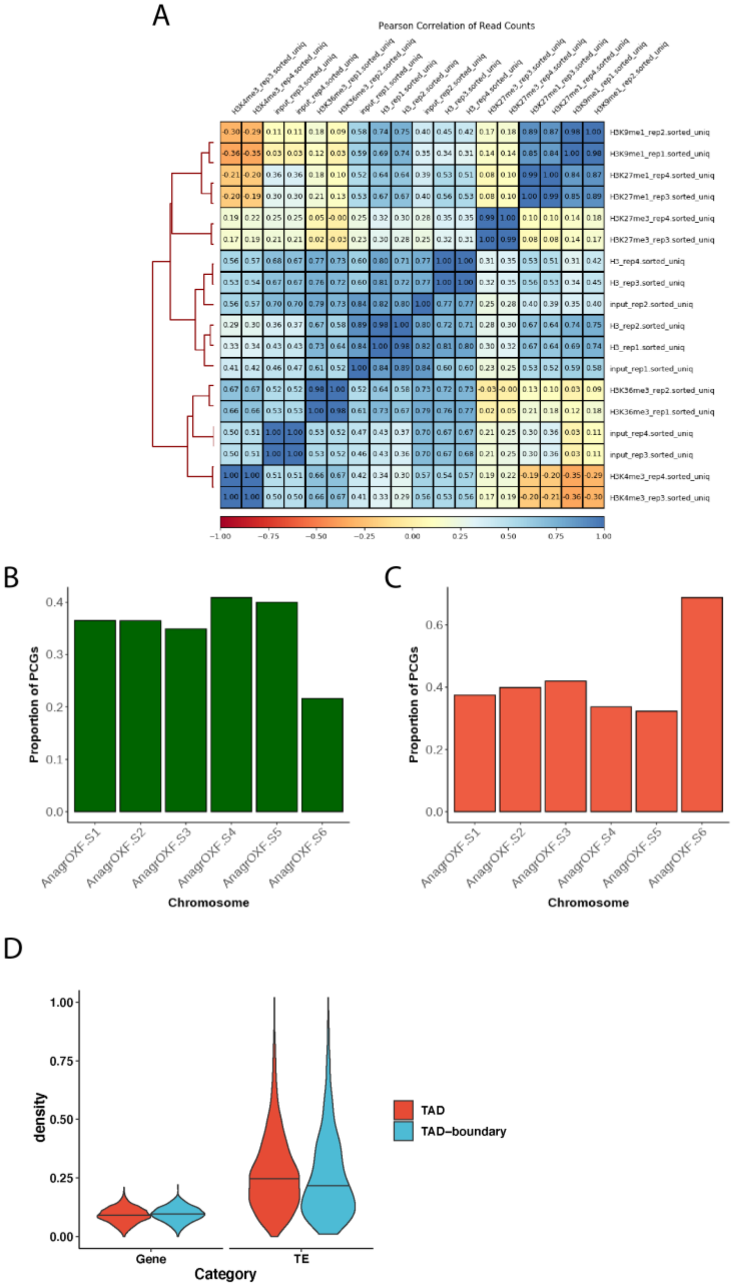


**Fig. S1. Quality control of ChIP-seq**

(A) Pearson correlation matrix showing that biological replicates of each mark cluster together.

(B) and (C) Bar plot showing the proportion of PCGs (A) or TEs (B) in each chromosome. Proportions were calculated as the total length of the features in each chromosome is divided by the length of each chromosome.

(D) Violin plots showing density distribution of genomic features (PCGs or TEs) in TADs and TAD boundaries. The density of genomic features was calculated as the number of features in each 40 kb window, divided by window length, and plotted for TADs and TAD boundaries. The median value is represented by a solid horizontal line.

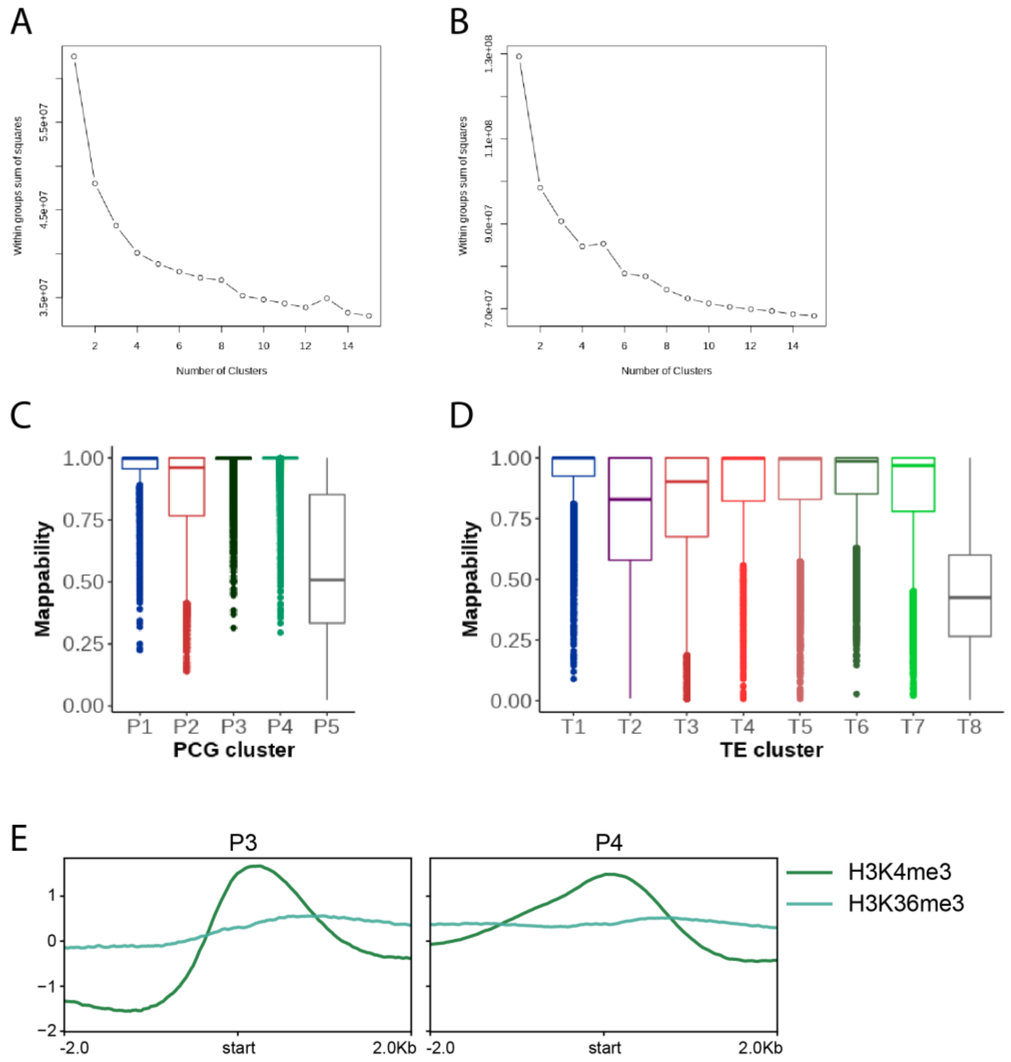


**Fig. S2. K-means clustering of ChIP-seq data over PCGs and TEs**

(A) and (B) Within groups sum of squares calculated using the output files of computeMatrix command over PCGs (A or TEs (B) are plotted against numbers of clusters.

(C) and (D) boxplots indicating the mappability of PCGs (C) and TEs (C) per cluster.

(E) Profile plot showing log2 ChIP/H3 enrichment of H3K4me3 and H3K36me3 over PCGs in clusters P3 and P4. Sequences 2 kb upstream and downstream of the start codon are included.

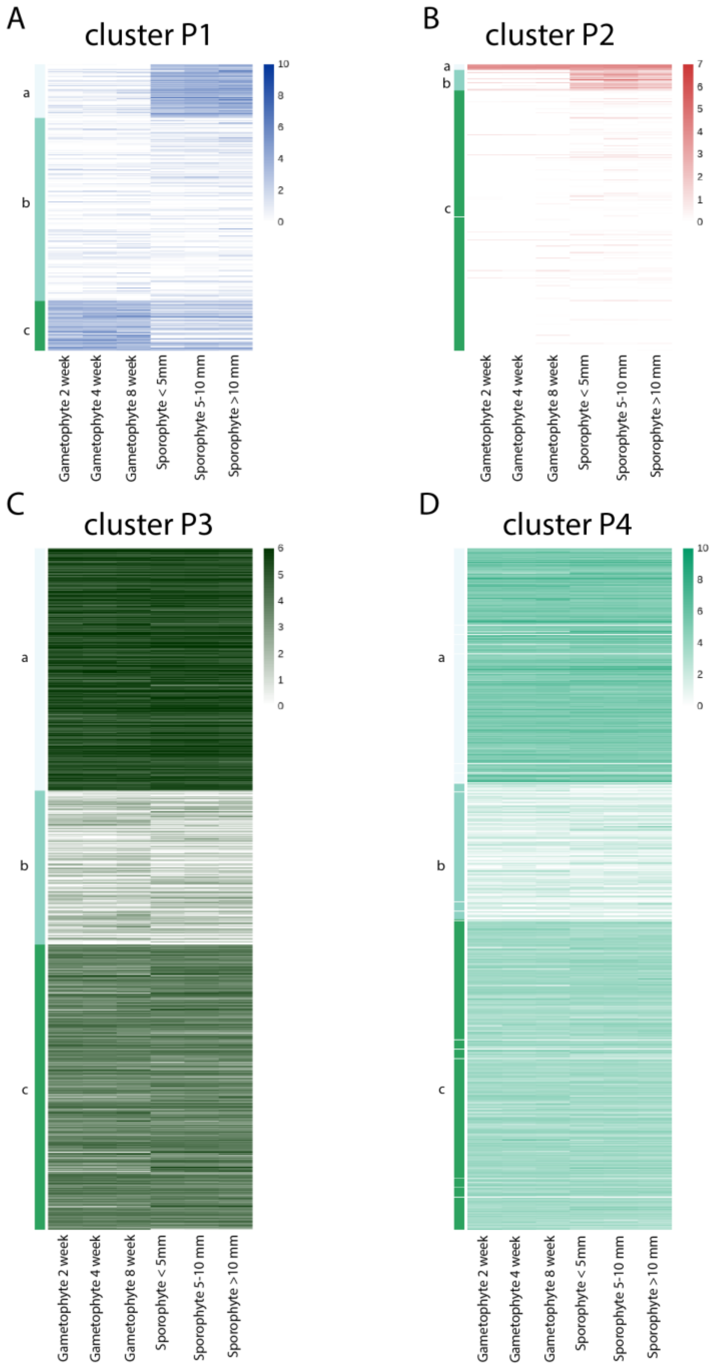


**Fig. S3. Expression level of PCGs in gametophyte and sporophyte**
(A), (B), (C) and (D) Heatmaps showing expression levels of PCGs in gametophyte and sporophyte tissue per cluster. Expression levels are indicated by arcsine TPM.

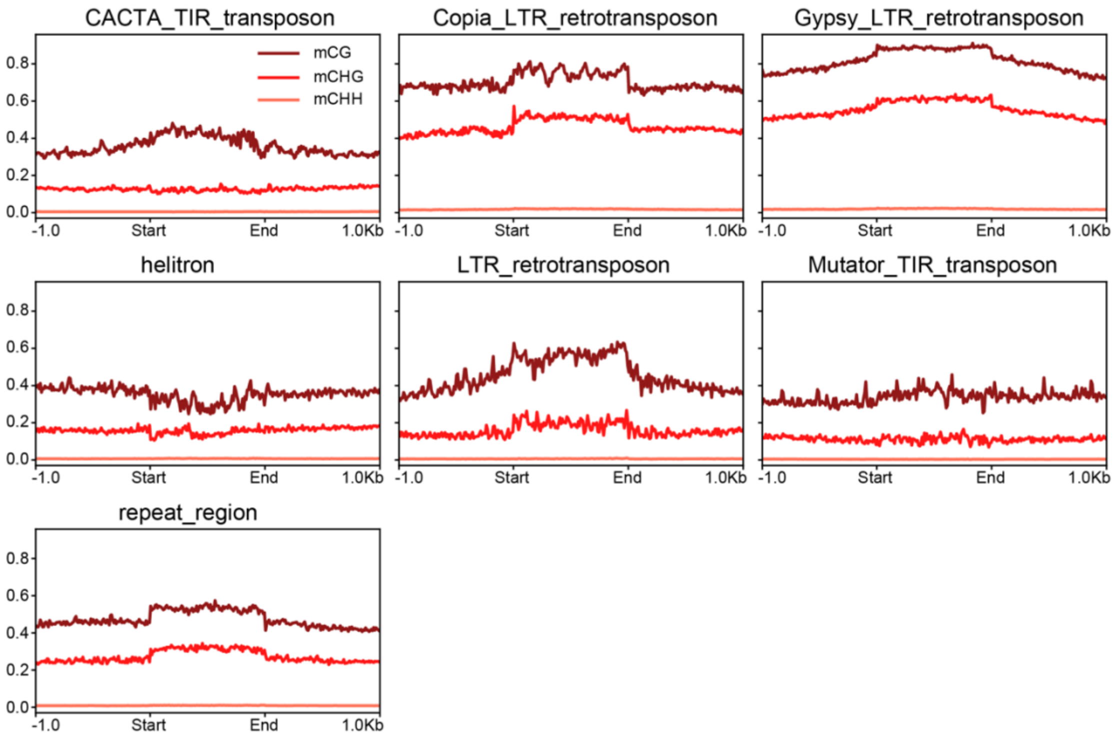


**Fig. S4. DNA methylation levels over each TE family**
Profile plot of CG, CHG, and CHH methylation levels over TEs per TE family. Each TE annotation is scaled to 1 kb and sequences 1 kb upstream and downstream are included. Average methylation over 10 bp bins is plotted.

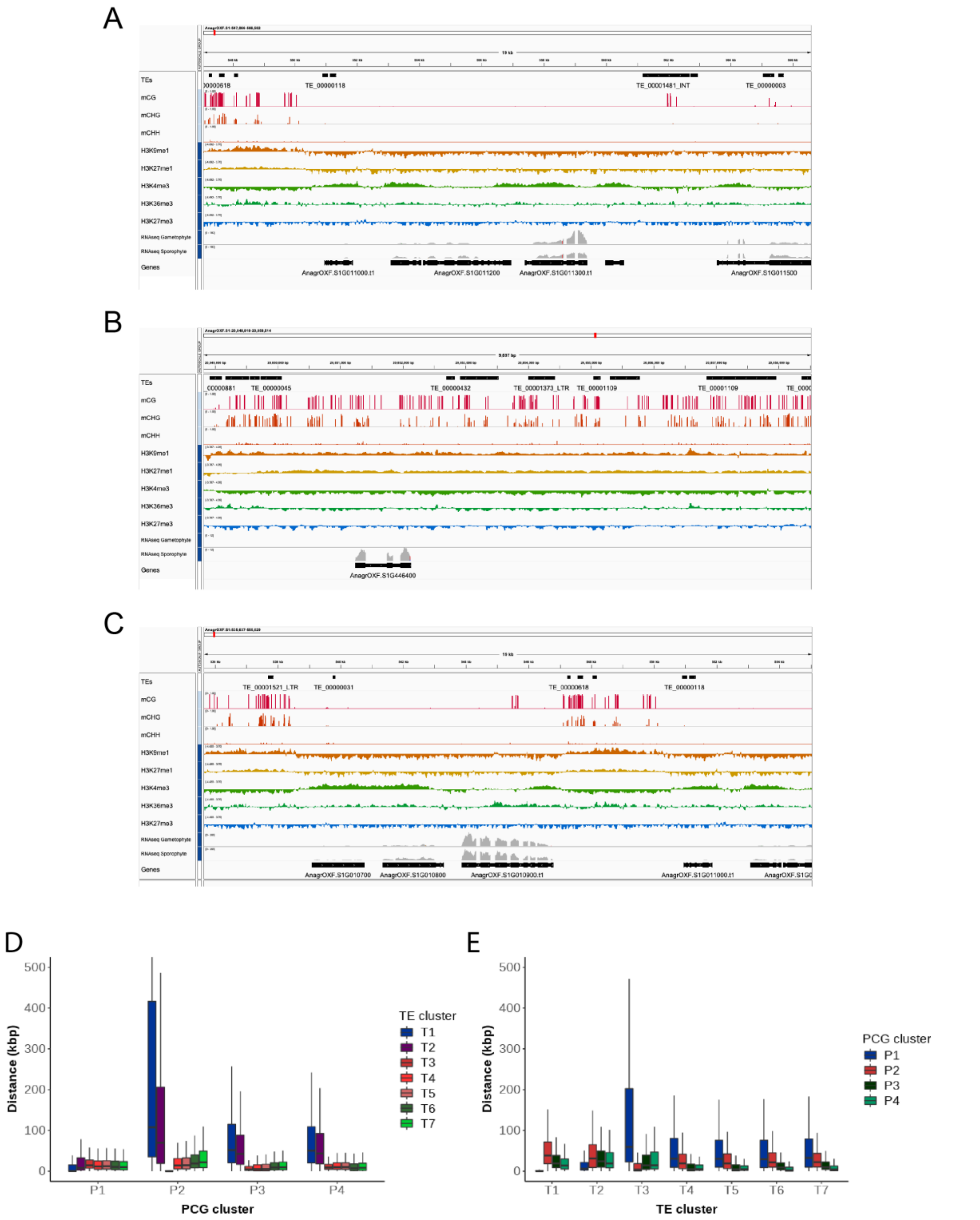


**Fig. S5. Distances between PCGs and TEs per cluster**

(A), (B) and (C) Integrative Genomics Viewer (IGV) browser screenshot demonstrating PCGs in the cluster P4 forming a small euchromatic island (A), PCGs in the cluster P2 surrounded by TEs (B) and TEs in the promoter region of PCGs in the cluster P3. The regions shown are 19 kb in length in (A) and (C) or 9.5 kb in (B) from the scaffold AnagrOXF.S1. PTM tracks are bigwig files scaled by H3 coverage in 10-bp windows. DNA methylation tracks are bigwig files showing methylation levels of each cytosine site covered by at least 10 reads. ‘‘TEs’’ and ‘‘Genes’’ tracks are annotation files for TEs and genes, respectively. ‘‘RNA-seq’’ tracks are bigwigs of mapped RNA-seq reads from gametophyte tissue and sporophyte tissue [39]. Scales are noted in square brackets in each track.

(D) Boxplots of distances between each gene and the nearest TE per TE cluster. Briefly, each gene is compared to all TEs belonging to a TE cluster to find its nearest neighbor. Genes are divided based on the gene cluster they belong to. Distances are in kilobase pairs (kbp). Colored boxes represent interquartile range, and lines represent median values. Outliers are not shown.

(E) Boxplot of distances between each TEs and the nearest PCG per PCG cluster. Briefly, each TE is compared to all PCGs belonging to a PCG cluster to find its nearest neighbor. TEs are divided based on the TE cluster they belong to. Distances are in kilobase pairs (kbp). Colored boxes represent interquartile range, and lines represent median values. Outliers are not shown.
